# Cross-species regulatory sequence activity prediction

**DOI:** 10.1101/660563

**Authors:** David R. Kelley

## Abstract

Machine learning algorithms trained to predict the regulatory activity of nucleic acid sequences have revealed principles of gene regulation and guided genetic variation analysis. While the human genome has been extensively annotated and studied, model organisms have been less explored. Model organism genomes offer both additional training sequences and unique annotations describing tissue and cell states unavailable in humans. Here, we develop a strategy to train deep convolutional neural networks simultaneously on multiple genomes and apply it to learn sequence predictors for large compendia of human and mouse data. Training on both genomes improves gene expression prediction accuracy on held out sequences. We further demonstrate a novel and powerful transfer learning approach to use mouse regulatory models to analyze human genetic variants associated with molecular phenotypes and disease. Together these techniques unleash thousands of non-human epigenetic and transcriptional profiles toward more effective investigation of how gene regulation affects human disease.

## Introduction

Predicting the behavior of any nucleic acid sequence in any environment is a primary objective of gene regulation research. In recent years, machine learning approaches to directly tackle this problem have achieved significant accuracy gains predicting transcription factor (TF) binding, chromatin features, and gene expression from input DNA sequence (1–7). These models have been applied to study genetic variation in populations and generate mechanistic hypotheses for how noncoding variants associated with human disease exert their influence (3–8). Estimates for how mutations influence regulatory activity have also revealed insights into regulatory evolution and the robustness of genes to such mutations (7).

Deep convolutional neural networks have achieved state of the art performance for many regulatory sequence activity prediction tasks in humans and other species (2, 4–7). The complexity of mammalian gene regulation and these models’ impressive but imperfect predictions suggest room for improvement remains. In particular, distal regulation by enhancers is incompletely captured by existing models, which do not attempt to consider sequence beyond ∼20 kb(5, 7). Obtaining more training data is a reliable strategy to improve model accuracy. The research field continues to generate new functional genomics profiles, but these merely deliver additional labels for the existing sequence data; fitting more expressive and accurate models would benefit more from entirely new training sequences. Individual human genomes differ only slightly from each other, so acquiring functional profiles for more humans is unlikely to provide this boost. Artificially designed sequences can offer more data for specific tasks, but only short sequences can be effectively manipulated and their profiling is limited to cell lines that cannot represent the full complexity of human tissues (9–13).

Non-human species offer a potential source of this desired additional training data. Regulatory sequence evolves rapidly, but TF binding preferences are highly conserved due to the drastic effect that modifying affinity for many thousands of binding sites would confer on the organism (14–16). Thus, we hypothesized that regulatory programs across related species have enough in common that jointly training on data from multiple species will improve regulatory sequence activity models derived by machine learning. To demonstrate the concept, we chose the mouse as a distant mammal with substantial functional genomics data available (17).

In addition to serving as a source of more genomic sequence, mouse experiments can explore biological states that are challenging or unethical to acquire in humans, e.g. profiling mouse development, disease, and genome modifications. If context-specific regulatory programs are sufficiently conserved across species, then models trained to predict these mouse data may be applicable to impute human genome profiles to study human regulatory sequences and genetic variation. Although models trained on mouse will not achieve the performance of analogous human models, they may serve as a useful approximation and produce novel variant annotations when the human data are unavailable.

In this work, we trained a deep convolutional neural network to jointly learn the complex regulatory programs that determine TF binding, DNA accessibility, and transcription using the ENCODE and FANTOM compendia of thousands of functional genomics profiles from hundreds of human and mouse cell types. We benchmarked single versus joint training and found that jointly training on human and mouse data leads to more accurate models for both species, particularly for predicting CAGE RNA abundance. We demonstrated that mouse regulatory programs can be transferred across species to human where they continue to make accurate tissue-specific predictions. Applying this procedure to predict human genetic variant effects revealed significant correspondence with eQTL statistics and proved insightful for studying human disease.

## Results

### Multi-genome training improves gene expression prediction accuracy

We applied the Basenji software and framework to predict functional genomics signal tracks from only DNA sequence (5). The neural network takes as input a 131,072(= 2^17^) bp sequence, transforms its representation with iterated convolution layers, and makes predictions in 128 bp windows across the sequence for the normalized signal derived from many datasets (Fig 1, Methods). We applied an architecture that uses residual connections to alleviate the strain of vanishing gradients in deep network optimization to improve generalization accuracy (Figure S1) (18). Training on multiple genomes required several further developments (Methods). Most importantly, we modified the train/valid/test split of the genomic sequences to ensure that homologous regions from different genomes did not cross splits (Methods); without this extra care, we might overestimate generalization accuracy.

**Figure 1:**
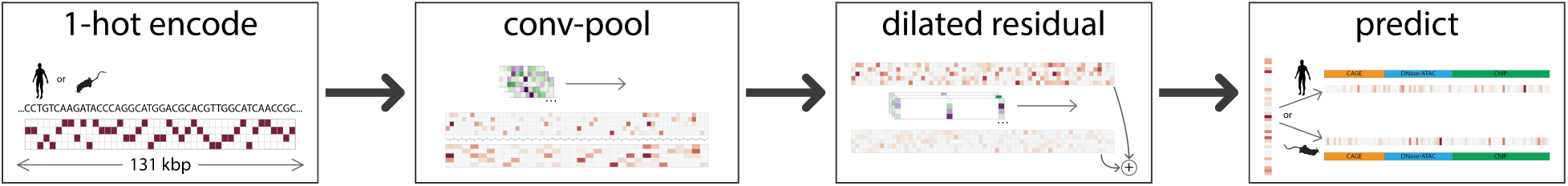
Predicting regulatory sequence activity for human and mouse genomes. We predict the regulatory activity of DNA sequences for multiple genomes in several stages (Methods). The model takes in 131,072 bp DNA sequences, encoded as a binary matrix of four rows representing the four nucleotides. We transform this representation with seven iterated blocks of convolution and max pooling adjacent positions to summarize the sequence information in 128 bp windows. Green and purple heatmaps represent convolution filter weights; red and white heatmaps represent pooled sequence vectors. To share information across the long sequence, we apply eleven dilated residual blocks, consisting of a dilated convolution with exponentially increasing dilation rate followed by addition back into the input representation. Finally, we apply a linear transform to predict thousands of regulatory activity signal tracks for either human or mouse. All parameters are shared across species except for the final layer.

We assembled training data consisting of 6,956 human and mouse quantitative sequencing assay signal tracks from the ENCODE and FANTOM consortiums (Methods). These data describe regulatory activity across tissues and isolated cell types using several techniques—DNase and ATAC-seq to measure DNA accessibility, which typically mark TF-bound sites, and ChIP-seq to map TF binding sites and histone modification presence (19, 20). The FANTOM data consists of RNA abundance profiling with CAGE, where the 5’ end of the transcript is sequenced (21). These 5’ RNA profiles are independent of splicing and allow us to provide DNA sequence without gene annotations, which would not be the case for RNA-seq (5). In addition, we added several mouse datasets describing cell states that are unavailable for humans: (1) a single cell ATAC-seq atlas from 13 tissues clustered to 85 distinct profiles (22) and (2) several TF and chromatin profiles obtained over 24 hour time courses in the liver to study circadian rhythms (S1 Table).

To measure the influence of multi-genome training on generalization accuracy, we trained three separate models on these data: one jointly fit to both human and mouse, one to human data alone, and one to mouse data alone. For each scenario, we fit the same model architecture and hyperparameters. We trained the models to minimize log Poisson loss and assessed accuracy by computing Pearson correlation between the predictions and observed data for each dataset.

The joint training procedure improved test set accuracy for 94% of human CAGE and 98% of mouse CAGE datasets (binomial test p-values 1e-16 and 1e-16), increasing the average correlation by .013 and .026 for human and mouse respectively (Fig 2a,c). For DNase, ATAC, and ChIP, joint training improved predictions by a lesser margin relative to single genome training; average test set correlation increased for 55% of human and and 96% of mouse datasets (binomial test p-values 3e-11 and 1e-16) (Fig 2b,d). The datasets where single genome accuracy exceeded joint tended to be consistent across a second independent batch of training runs (Figure S3). ChIP-seq experiments in the cancer cell lines K562 and MCF-7 were significantly enriched in this set, suggesting that modeling these datasets may slightly suffer due to absence of an analogue across species or somatic mutation divergence from the reference genome.

**Figure 2:**
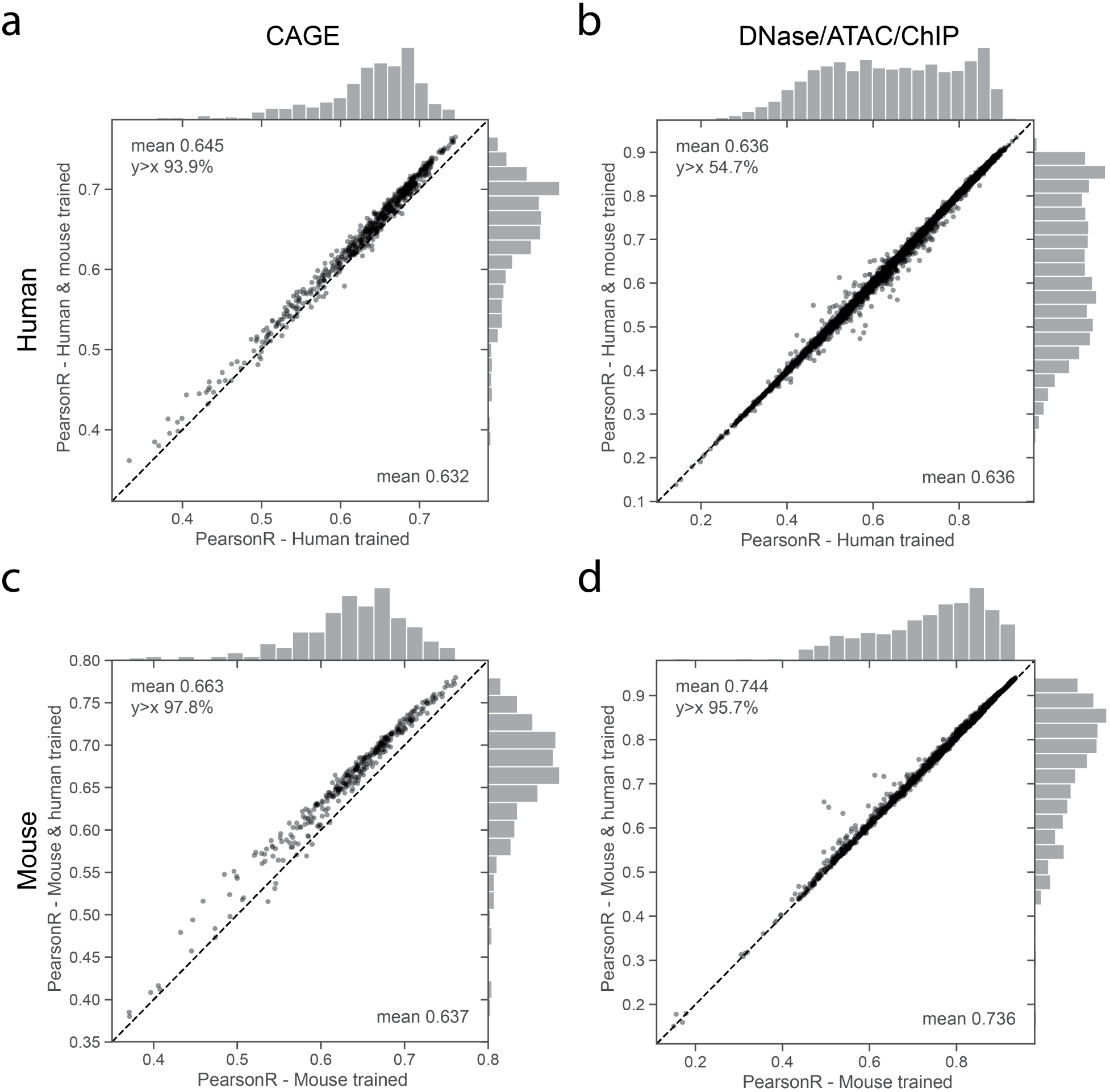
Training on human and mouse data improves generalization accuracy. We trained three separate models with the same architecture on human data alone, mouse data alone, and both human and mouse data jointly. For each model, we computed the Pearson correlation of test set predictions and observed experimental data for thousands of datasets from various experiment types. Points in the scatter plots represent individual datasets, with single genome training accuracy on the x-axis and joint training accuracy on the y-axis. For CAGE, training on multiple genomes increases test set accuracy on nearly all datasets for both human and mouse. For DNase/ATAC/ChIP-seq, test set accuracy improves by a smaller average margin.

CAGE has several properties that may explain the observed extra benefit of having more training data from multiple genomes. CAGE signal has a larger dynamic range than the other data, spanning orders of magnitude, fewer relevant sites in the genome, and more sophisticated transcriptional regulatory mechanisms that often involve distant sequences. Measured by gradient-based saliency analysis, the jointly trained model makes greater use of long range activating elements (*>* 10 kb) to predict CAGE signal at transcription start sites (TSSs) (Figure S2). Altogether, these results demonstrate that regulatory programs are sufficiently similar across the 90 million years of independent evolution separating human and mouse so that their annotated genomic sequences provide informative multi-task training data for building predictive models for both species.

### Regulatory sequence activity models transfer across species

Regulatory program conservation across related species has been observed in genome-wide functional profiles of TF binding and histone modifications (14–16). In matched tissue samples, similar TFs are typically present and those TFs have highly conserved motif preferences (16, 23). These findings suggest that a regulatory sequence activity model trained to predict for one species will also make usefully accurate predictions for matched samples from the other. This was recently demonstrated for enhancer annotations and histone modifications across a variety of mammals (24). To quantify this phenomenon for our models and data, we selected several diverse and representative tissues and cell types for which we could unambiguously match across species—cerebellum, liver, and CD4+ T cells. We extracted CAGE gene expression measurements from the TSSs for all human genes outside the training set and computed predictions for human and mouse versions of these tissues and cell types (Fig 3a). For this exercise, and those to follow, we used the jointly trained multi-task model and sliced out predictions of interest, but results were consistent for the model trained only on mouse data (Figure S4).

**Figure 3:**
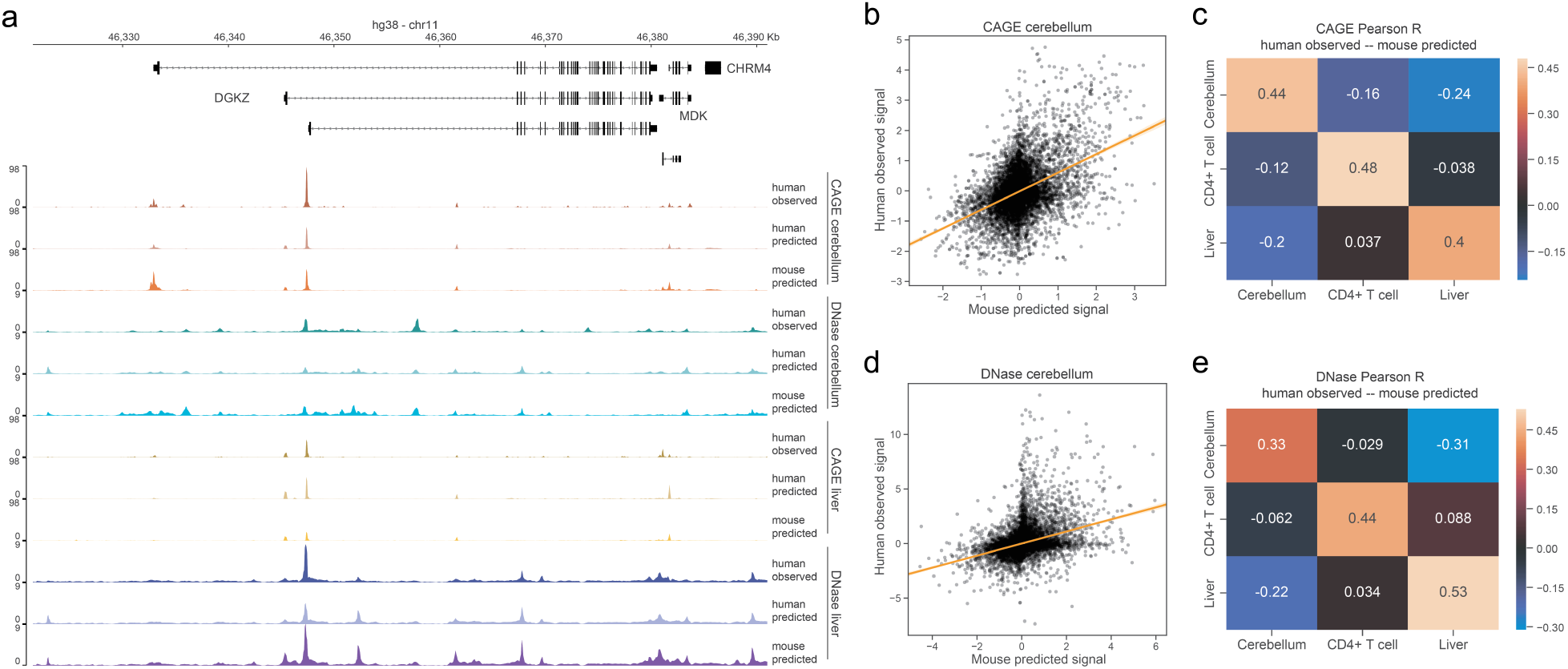
Regulatory programs are largely conserved across species. (a) Tissue-specific regulatory programs can be learned and transferred across species, exemplified here by CAGE and DNase data and predictions for cerebellum and liver. The “human predicted” tracks describe predictions for the human datasets displayed as “human observed”; “mouse predicted” tracks describe predictions for the matched mouse dataset. We scaled coverage tracks by their genome-wide means separately within all CAGE and all DNase/ATAC data. (b) Mouse predictions for cerebellum CAGE and (d) DNase correlate strongly with human data. For CAGE, points represent the top 50% most variable TSSs, where data or predictions were quantile normalized to align sample distributions, log transformed, and mean-normalized across samples. For DNase, points represent the top 10% most variable genomic sites (less than CAGE because we consider the whole genome rather than TSSs), where data or predictions were similarly quantile normalized to align sample distributions and mean-normalized across samples. The statistical trends were robust to most variable threshold choice. Scatter plot lines represent ordinary least squares regressions. (c,e) These correlations are specific to brain regions and not shared by other tissues, such as CD4+ T cells or liver.

Across human gene TSSs, we observed cross-species prediction accuracy of 0.73 Pearson correlation for mouse predictions to human observed signal averaged across these samples. This approaches the 0.75 correlation for human predictions to human observed signal, but does suggest that some genome-specific activity exists (Figure S5). To assess whether the model further captures and transfers tissue specificity, we normalized each TSS’s data or predictions by its mean across all CAGE datasets. Mean normalization removes correlation driven by accurate prediction of global cross-tissue activity. On this more challenging task, normalized mouse predictions achieved mean 0.40 Pearson correlation with normalized human data for the matched samples (Fig 3b,c). In contrast, normalized predictions compared to data from distinct tissues/cell types resulted in negative correlations (Fig 3c). Thus, the models have learned tissue and cell type specificity beyond a baseline level and are able to transfer that knowledge across species.

We repeated these analyses with DNase accessibility profiles for the same tissues and cell types to assess how general this transferability is for different data. Because most sites lack activity, we selected the top 10% most variable. We observed the same statistical trends for accessibility—high correlation between mouse predictions and human data for matched samples (mean 0.84) and specificity for scaled comparisons (Fig 3d,e).

### Mouse-trained models elucidate human genetic variant effects

A driving goal of regulatory sequence modeling is to predict the effect of human genetic variants on gene expression and downstream phenotypes. For any biallelic variant, we can predict signal across the surrounding genomic sequence for each allele and derive a summary score for the variant effect (Fig 4a). Here, we sum the signal across the sequence and take the difference between alleles. We can compute this score for every dataset using two forward passes of the convolutional neural network.

**Figure 4:**
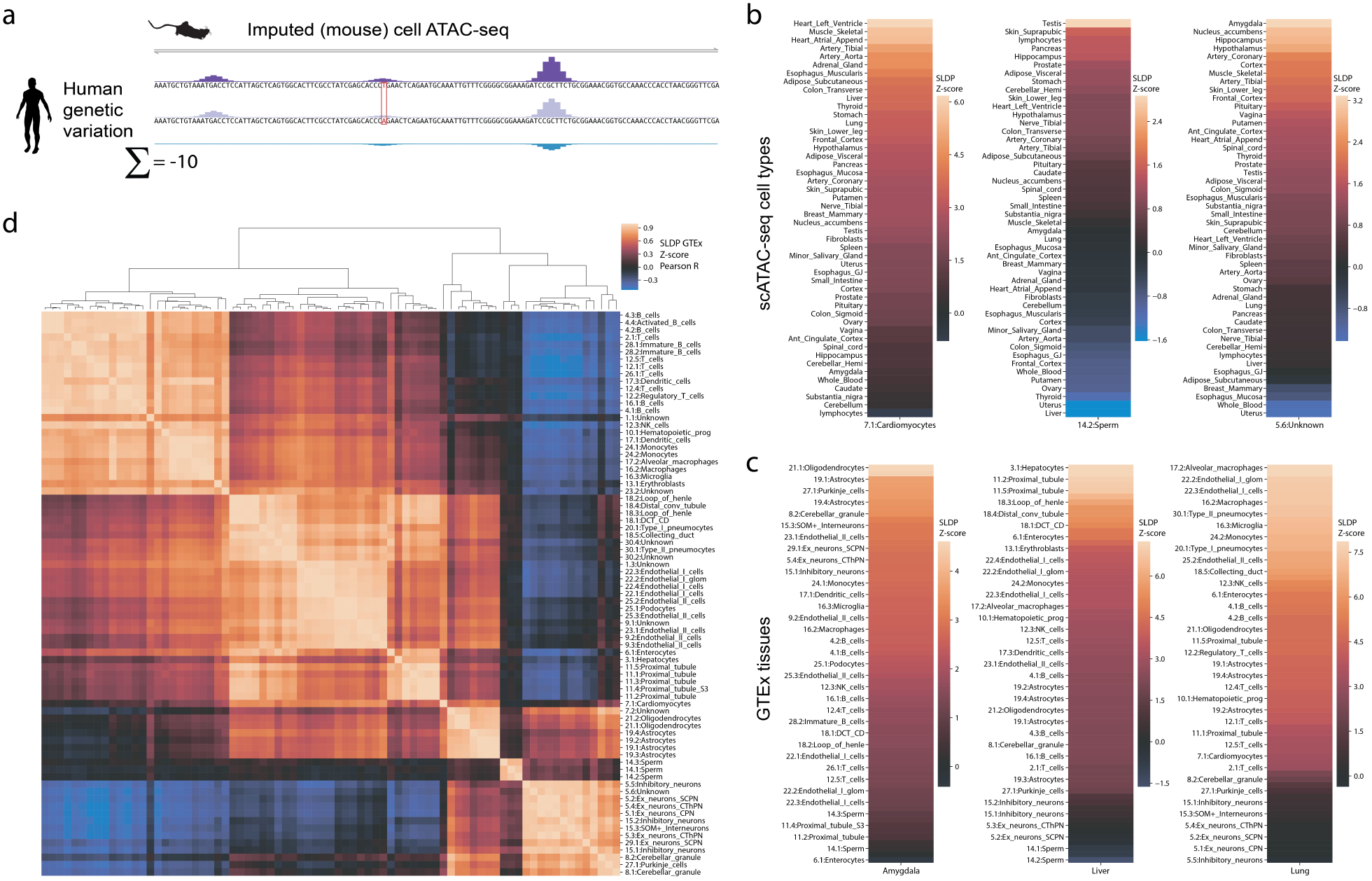
Mouse cell type accessibility predictions show a strong and specific statistical relationship with human eQTLs. (a) We predicted the effect of human genetic variants on imputed regulatory signal trained on mouse single cell ATAC-seq (scATAC) cluster profiles. We scored variants by subtracting the signal from the minor allele from that of the major and summing across the sequence. (b) We used signed linkage disequilibrium profile (SLDP) regression to compare the cell type-specific variant effect predictions to tissue-specific eQTL summary statistics from GTEx. Cell type profiles correspond best with the expected tissues. (c) GTEx tissues correspond best with the expected cell types. (d) Clustering scATAC cell types by their z-scores across GTEx tissues reveals the expected structure.

Models trained on mouse data allow one to predict the difference between how two human alleles would behave if they were present in the regulatory environment of mouse cells. Given the evidence that analogous human and mouse cells largely share regulatory programs, we hypothesized that models trained on mouse data would be insightful towards understanding human regulatory variants’ function. To test this hypothesis, we studied the Gene-Tissue Expression (GTEx) release v7a data of genotypes and gene expression profiles for hundreds of humans across dozens of tissues (25). In previous work, we showed that variant scores derived from Basenji predictions corresponded significantly with GTEx summary statistics (5). Here, we conducted a similar analysis using signed linkage disequilibrium profile (SLDP) regression to measure the statistical concordance between signed variant effect predictions and GTEx summary statistics (Methods) (8). SLDP distributes a signed annotation (i.e. our scores) according to a given population’s LD structure and compares it to a set of summary statistics. Using a permutation scheme, the method produces a signed Z-score that specifies the direction and magnitude of the relationship and a p-value describing its significance.

We focused on a dataset unique to the mouse—a single cell ATAC-seq atlas from 13 adult mouse tissues (22) that reported 85 distinct cell types after clustering analysis. Merging aligned reads for each cell type cluster produced pseudo-bulk coverage profiles, which we trained to predict. We sliced predictions for these datasets from the model trained jointly on all human and mouse data. We first asked whether coverage tracks derived from clustering single cell assays are amenable to Basenji modeling. Predictions for held out sequences achieved Pearson correlation ranging from 0.43-0.84 in 128 bp windows for these 85 profiles, which is in line with predictions for bulk DNase/ATAC-seq.

Human variant predictions for these models generally exhibited a strong, positive effect on GTEx summary statistics, in line with prior observations that increased accessibility typically increases gene expression. Furthermore, cell type predictions aligned well with anatomical expectations. For example, variant predictions for cardiomyocytes have the strongest correlation with GTEx measurements in the heart and skeletal muscle (Fig 4b). From the opposite direction, GTEx measurements for the liver have the strongest correlation with variant predictions for hepatocytes (Fig 4c). These results further support the claim that human and mouse cells share relevant regulatory factors and that our procedure can project these factors across species from mouse experiments to human variants.

For each pair of mouse ATAC cell types, we computed the correlation between their SLDP Z-scores across GTEx tissues (Fig 4d). The correlations revealed expected structure, with clusters representing the blood, endothelial cells, neurons, among others. The original authors abstained from annotating 9 of the 85 clusters. Through this procedure, we can suggest high-level annotations for 6/9 of the unknown clusters. For example, 5.6 appears similar to various neuron subtypes due to the strong statistical relationship between variant predictions and the GTEx brain tissue summary statistics (Fig 4b,d).

Next, we asked whether these or any other mouse datasets were informative above and beyond available human datasets. Theoretically, we can add scores for every human dataset to the SLDP background model, forcing the statistical test for mouse dataset scores to consider only the residual variance in GTEx summary statistics (Methods). We implemented a close approximation where we added 64 principal components of the variant by score matrix for human datasets, which explained 99.9% of the variance for CAGE and 99.3% for DNase/ATAC.

Even considering the human data, many mouse datasets still emerged as delivering orthogonal value by SLDP (Figure S6). Among CAGE data, developmental heart profiles from neonate and embryo stages had significant positive correlations with GTEx tibial artery (10 datasets with FDR *q <* 0.05) and left ventricle (41 datasets with FDR *q <* 0.05). These suggest that human adult heart gene expression depends on genetic variation that acts more prominently during early development, which is recovered more effectively by current data from mice. Among DNase/ATAC, variant predictions for the single cell hepatocytes above and a 24 hour time course to profile circadian rhythms of genome accessibility in the liver (26) showed a significant positive relationship with the GTEx liver statistics (7 datasets with FDR *q <* 0.05). GTEx samples were collected at a variety of day times, and previous work demonstrates that they can be ordered to recover circadian cycling genes (27). Our result suggests that available human DNase profiles fail to recover this variance, but uniquely obtainable mouse time series do.

Finally, we hypothesized that the improved accuracy of the jointly trained models on held out chromosome sequences would carry over to more effective variant analysis. To assess this, we computed SLDP Z-scores to GTEx tissues for all human and mouse CAGE and DNase datasets using models trained jointly or on single genomes. For both species, the jointly trained models achieved greater SLDP Z-scores than their single genome counterparts (Figure S7). Thus, multi-genome training leads to greater correlation between variant effect predictions and GTEx summary statistics.

### Mouse-trained models highlight mutations relevant to human neurodevelopmental disease

Having established the relevance and specificity of mouse dataset predictions for expression phenotypes, we asked whether these data could provide insight into the genetic basis of human disease. Mouse data has proven valuable for studying human genetic variants in previous work (17, 22), but these analyses were limited to studying variants in homologous sequences in their mouse genome context. Given the substantial regulatory sequence turnover between these genomes, this limitation is severe. The predictive framework here avoids this limitation by mapping the learned mouse regulatory program to the human genome setting for all variants.

To explore the utility of this procedure for studying human disease, we retrieved a recent dataset of 1902 quartet families from the Simons Simplex Collection (28) with whole genome sequencing of a mother, father, child affected by autism, and unaffected sibling. In these data, the offspring have an average of 67 de novo mutations, which have a slight enrichment in promoters (29). Recent work demonstrated that variant effect predictions further differentiate autism cases from their unaffected sibling controls (30). We hypothesized that predictions using models trained on mouse data would also distinguish the disease and perhaps provide additional insight via novel developmental profiles.

We applied the model to predict how each de novo mutation would influence signal in 357 mouse CAGE profiles of tissues and cell types throughout the body. Mann-Whitney U (MWU) tests revealed significantly more negative predictions in the case versus control variant sets for 246 CAGE profiles at FDR *<* 0.1 (Fig 5a). Appreciating the correlations in these data, we also transformed the variants by predictions matrix with PCA to represent each variant by its first principal component score (which explained 51% of the variance). In principal component space, the MWU test comparing case and control variants was significant with p-value Most leading datasets described brain regions and cell types; the 76 brain dataset p-values were less than non-brain data with p-value 1 *×*10^−10^ by MWU test. Human CAGE predictions validated the case set enrichment of negative predictions, particularly for brain-specific tissues (Figure S8).

**Figure 5:**
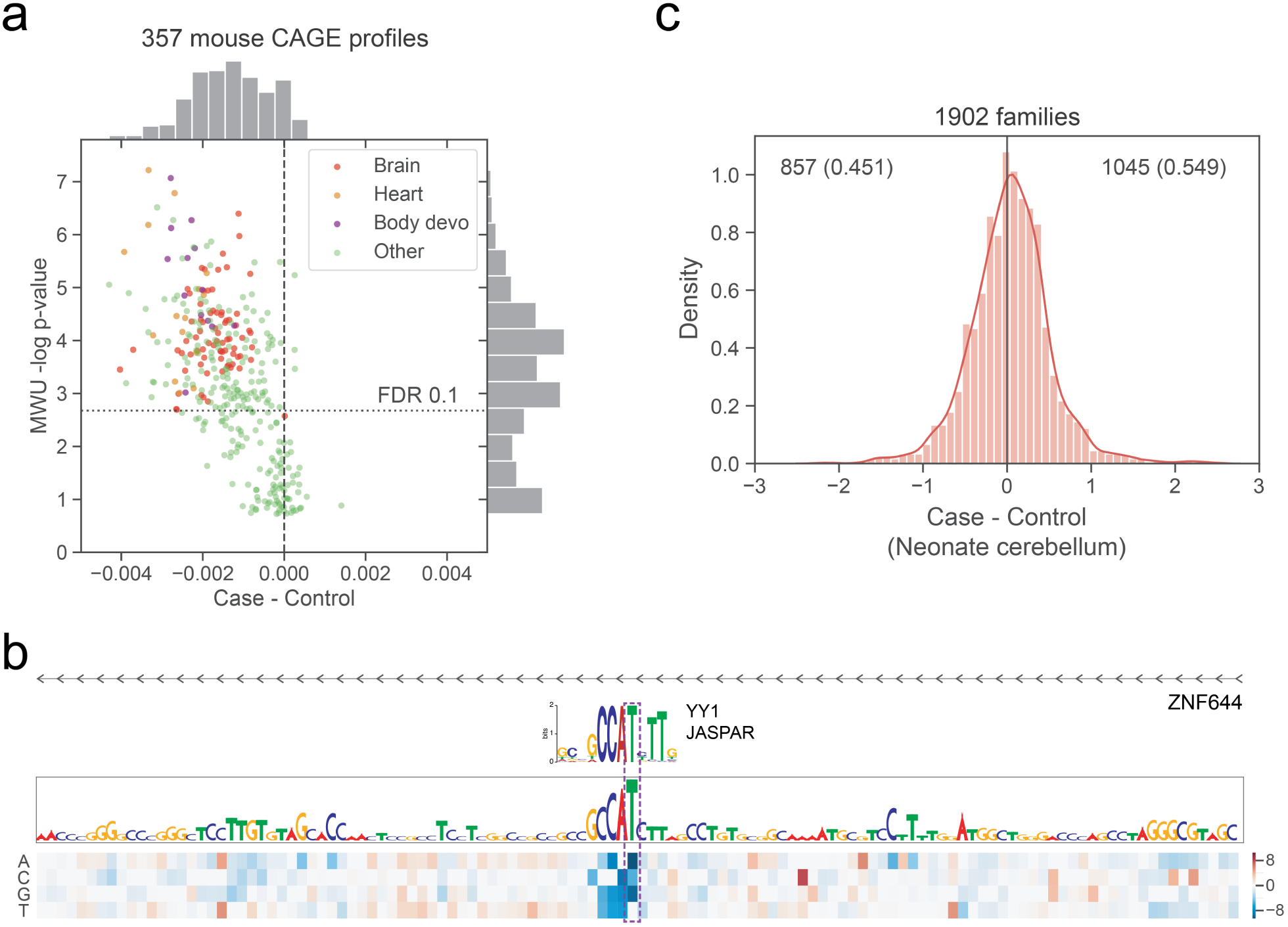
Human de novo variant predictions for mouse data enrich for autism cases versus controls. (a) We predicted the influence of 234k de novo variants split between cases and controls on 357 CAGE datasets in mouse. For each dataset, we computed a Mann-Whitney U (MWU) test between case and control sets and corrected for multiple hypotheses using the Benjamini-Hochberg procedure. Predictions for many datasets were enriched for more negative values in the cases, driven largely by brain, heart, and whole body developmental profiles. Each dataset’s x-axis position is the mean inverse hyperbolic sine over case variants minus the equivalent over control variants. The mean inverse hyperbolic sine transformation is similar to logarithm, but gracefully performs the symmetric transformation for negative values. (b) A case variant at chr1:91021795 modifies a critical T in a YY1 motif to an A in the promoter region of *ZNF644*. (c) At the individual level, a simple score summing all negative predictions for the leading dataset describing neonate cerebellum significantly separates cases from their matched controls.

Highly negative predictions indicate mutations that disturb active regulatory elements. For example, a case variant upstream of *ZNF644* modifies a critical nucleotide in a consensus motif for the transcription factor YY1, which the model identifies as active and relevant (Fig 5b). *ZNF644* has considerable evidence for intolerance to loss of function mutations in the Genome Aggregation Database v2.1.1 (gnomAD) with probability 0.999 of intolerance (31). YY1 has been implicated in processes that determine the three-dimensional positioning of promoters and enhancers (32). Thus, we hypothesize that the variant modifies the enhancer regulation of this critical protein.

Perhaps unexpectedly, 15 datasets describing the developing heart also emerged from this analysis (Fig 5a). This result is supported by whole genome sequencing of congenital heart disease probands, which has revealed affected gene sets that overlap significantly with those observed in neurodevelopmental sequencing efforts like this one (33, 34). In addition to the brain and heart, whole body profiles from the embryo and neonate stages also have p-values among the lowest.

This significant enrichment indicates that variant effect predictions may help classify disease at the individual level. For each individual, we computed a simple risk score by summing the negative predictions in the neonate cerebellum dataset. This score suggests more deleterious de novo variants for 54.9% of the cases versus their controls (binomial test p-value 9 *×* 10^−6^) (Fig 5c). Thus, this approach is a strong candidate for inclusion with complementary feature sources from coding mutations and structural variation to continue to characterize this incompletely understood disorder.

## Discussion

In this work, we developed a novel convolutional neural network architecture and multiple species training procedure to enable one model to train on 6956 functional genomics signal tracks annotating the human or mouse genomes. We observed that training jointly on both species produced models that make more accurate predictions on unseen test sequences relative to models trained on a single species. Regulatory sequence activity predictions for human sequences in mouse tissues correlate well with datasets describing the corresponding human tissues. Model predictions for altered regulatory activity of human genetic variants made with respect to mouse datasets have a strong statistical concordance with tissue-specific human eQTL measurements. Mouse machine learning models can be used to study human disease, exemplified by enrichment of deleterious predictions among de novo autism variants relative to control sets.

We focused here on human and mouse because both species have been comprehensively studied with genome-wide functional genomics. Our observation that joint training on these two genomes improves prediction accuracy opens the possibility of more complex schemes for training on larger numbers of genomes. Given the substantial evolutionary distance between human and mouse, regulatory annotations for all mammalian genomes are likely to provide similarly useful training data. Primate genomes will be particularly interesting to explore; their tissues and cell types will more closely match those of human, but their sequences are far more similar. Prediction accuracy improved more for CAGE gene expression measurements than accessibility or ChIP-seq, which suggests that the number of events and their regulatory complexity are relevant features for determining whether multiple genome training will be worthwhile. Efforts to predict spatial contacts between chromosomes as mapped by Hi-C and its relatives likely fit this criteria, and we hypothesize that training sequence-based models on human and mouse data together will be fruitful.

Much prior work has revealed the similarity of regulatory programs across species, but transferring knowledge gleaned from an accessible model organism (such as mouse) to another of interest (such as human) has remained challenging. Existing approaches rely on whole genome alignments to transfer annotations from one genome to the other (22, 35). These approaches are constrained by the quality of the alignment, which is a notoriously challenging bioinformatics problem (36), and the limited proportion of each genome that aligns (40% for human and 45% for mouse). Here, we demonstrated an alternative approach where a machine learning model trained on the model organism data compresses the relevant knowledge into its parameters, which can then be applied to make predictions for sequences from the genome of interest. Substantial research in transfer learning with neural networks for natural language processing motivates and supports the viability of this procedure (e.g. (37)). The strong tissue-specific statistical relationship between human genetic variant predictions from model parameters trained to predict mouse annotations and GTEx tissue-specific eQTLs highlights the successful nucleotide resolution of our mouse to human transfer learning. The Gene Expression Omnibus (GEO) contains tens of thousands of mouse functional genomics profiles, many describing experiments impossible in humans. For example, we included dozens of datasets describing mouse liver profiles over 24 hour time courses to study the circadian rhythms of gene expression and chromatin. Models trained to predict all datasets, as well as open source software to compute these predictions and train new models on users’ own data, are available in the Basenji software package (38).

## Methods

### Functional genomics data

In this work, we studied quantitative sequencing assays performed on human and mouse samples. Specifically, we focused on DNase and ATAC-seq profiling DNA accessibility, ChIP-seq profiling TF binding or histone modifications, and CAGE profiling RNA abundance derived from 5’ transcription start sites. Preprocessing these data effectively is critical to successful machine learning. Our primary preprocessing objective is to denoise these data to the relevant signal at nucleotide-resolution.

We largely followed the preprocessing pipeline described in prior research introducing the Basenji framework (5)] The standard pipeline through which experimental data flowed follows:

1. Trim raw sequencing reads using fastp, which can automatically detect and remove unwanted adapter nucleotides (39).
2. Align reads using BWA to hg38 or mm10 and requesting 16 multi-mapping read positions (40).
3. Estimate nucleotide-resolution signal using an open source script (*bam_cov.py*) from the Basenji software that distributes multi-mapping reads, normalizes for GC bias, and smooths across positions using a Gaussian filter (5).

However, we varied from this standard pipeline for all data available from the ENCODE consortium website, which is 4,506 human and 1,019 mouse experiments. These data have been thoughtfully processed using open source pipelines and are available for download at several stages, including log fold change signal tracks in BigWig format (41). Rather than reprocess these data without full knowledge of how replicate and control experiments match, we chose to use these signal tracks directly. The Seattle Organismal Molecular Atlas (SOMA) server provides a single cell mouse ATAC-seq atlas (22). These data are also available in log fold change BigWig format, and we similarly chose to use these rather than reprocess the single cell data. We clipped negative values in all such BigWig tracks to zero.

We applied several transformations to these tracks to protect the training procedure from large incorrect values. First, we collected blacklist regions from ENCODE and added all RepeatMasker satellite and simple repeats (42), which we found to frequently collect large false positive signal (43). We further defined unmappable regions of *>*32 bp where 24-mers align to *>*10 genomic sites using Umap mappability tracks (44). We set signal values overlapping these regions to the 25^*th*^ percentile value of that dataset. Finally, we soft clipped high values with the function *f* (*x*) = *min*(*x, t*_*c*_ + *sqrt*(*max*(0, *x* − *t*_*c*_))). Above the threshold *t*_*c*_ (chosen separately for each experiment and source), this function includes only the square root of the residual *x* − *t*_*c*_ rather than the full difference.

When replicate experiments profiling the same or related samples were available, we averaged the signal tracks. Altogether, the training data includes 638 CAGE, 684 DNase/ATAC, and 3991 ChIP datasets in human and 357 CAGE, 228 DNase/ATAC, and 1058 ChIP datasets in mouse. S1 Table describes all data with preprocessing parameters. Code to preprocess typical functional genomics data formats into TensorFlow input formats is available from https://github.com/calico/basenji.

### Model architecture

We modeled genomic regulatory sequence activity signal as a function of solely DNA sequence using a convolutional neural network. Such deep learning architectures have excelled for many similar tasks (2, 4–6). We follow our prior work in analyzing large 131 kbp sequences in order to consider long range interactions.

The first stage of the architecture aims to extract the relevant sequence motifs from the DNA sequence using the following block of operations:

1. Convolution width 5 (or 15 in first layer)
2. Batch normalization
3. Gaussian Error Linear Unit (GELU) activation
4. Max pool width 2

We applied this block seven times so that each sequence position represents 128 bp, increasing the number of filters from an initial 288 by 1.1776x each block to 768 filters by the end. The GELU activation slightly outperformed the more common ReLU in our benchmarks (45).

The second stage of the architecture aims to spread information across the sequence to model long range interactions. In prior work, we applied densely connected dilated convolutions for this task (5). Here, we applied a related but more effective variation, which is related to a strategy applied for DNA sequence analysis in the SpliceAI system (46). Recent deep learning research has revealed that skip connections between layers where one layer’s representation is directly added to a subsequent layer’s representation relieve vanishing gradients and improve gradient descent training (18). Thus, we applied the following series of operations:

1. GELU activation
2. Dilated convolution width 3, dilation rate *d*, 384 filters
3. Batch normalization
4. GELU activation
5. Convolution width 1, back to 768 filters
6. Batch normalization
7. Dropout probability 0.3
8. Addition with the block input representation before step 1.

We applied this block eleven times, increasing the dilation rate *d* by 1.5x each time. Relative to the densely connected version, the dilated residual blocks lead to improved generalization accuracy (Figure S1).

In the final stage, we first transformed this 1024×768 (length × filters) representation of 128 bp windows with an additional width 1 convolution block using 1536 filters and dropout probability 0.05. To make predictions for either 5313 human or 1643 mouse datasets, we applied a final width one convolution followed by a softplus activation (*f* (*x*) = *log*(1 + *e*^*x*^)) to make all predictions positive. We attached a genome indicator bit to each sequence to determine which final layer to apply.

We trained to minimize a Poisson log likelihood in the center 896 windows, ignoring the far sides where context beyond the sequence is missing. The Poisson model is not technically appropriate for the log fold change tracks. However, by clipping negative values to zero, the distribution of values resembles that from our standard processing. Clipping to zero focuses attention on signal magnitude in regions where relevant signal is present and away from less relevant signal fluctuations in background regions. On a subset of data, we observed that using the log fold change track did not decrease accuracy or the utility of the model for genetic variant analysis.

We implemented the network in TensorFlow and used automatic differentiation to compute gradients via back propagation (47). We minimized with stochastic gradient descent (SGD) on batches of 4 sequences. We stopped training when the loss on a validation set had not improved for 30 epochs and returned to the model weights that had achieved the minimum validation loss. We performed several grid searches to choose model and optimization hyper parameters for the following sets: (1) SGD learning rate and momentum; (2) initial convolution filters and convolution filter multiplication rate; (3) dilated convolution filters and dropout rate; final convolution filters and dropout rate.

Data augmentation describes a suite of techniques to expand the implicit size of the training dataset from the perspective of model training by applying transformations that preserve annotations to data examples. We tiled the 131,072 bp sequences across the chromosomes by 65,599 bp, representing a 50% overlap minus 63 bp in order to also shift the 128 window boundaries and max pooling boundaries. During training, we cycled over combinations of two transformations that maintain the relationship between sequence and regulatory signal while changing the model input: (1) reverse complementing the sequence and reversing the signal; (2) shifting the sequence 1-3 bp left or right. Both transformations improved test accuracy and reduce overfitting in our benchmarks.

Model implementations and instructions for re-training, predicting, and modifying them are available from https://github.com/calico/basenji.

### Multi-genome training

Training on multiple genomes containing orthologous sequence complicates construction of holdout sets. Independently splitting each genome’s sequences would allow training on a human promoter and testing on its mouse orthologue. If the model memorized conserved elements of the sequence, rather than learning a general function, we might overestimate generalization accuracy.

We used the following procedure to minimize occurrence of this potential issue:

1. Divide each genome into 1 mb regions.
2. Construct a bipartite graph where vertexes represent these regions. Place edges between two regions if they have *>*100 kbp of aligning sequence in a whole genome alignment.
3. Find connected components in the bipartite graph.
4. Partition the connected components into training, validation, and test sets.

We used the hg38-mm10 syntenic net format alignment downloaded from the UCSC Genome Browser site (48). Using this procedure, we set aside approximately 12% of each genome into validation and test sets respectively. Stricter parameter settings created a single large connected component that did not allow for setting aside enough validation and test sequences.

Another complication of training on multiple genomes arises from imbalance between each genome’s sequences and datasets. We extracted 38.2k human and 33.5k mouse sequences for analysis. We assembled batches of sequences from one genome or the other, chosen randomly proportional to the number of sequences from each genome. The overall loss function comprises a term for every target dataset summed, which leads to larger step magnitudes for batches of human sequences that are annotated with *>*3 times more datasets. Explicit weighting could be applied to preference training towards a particular species, but we found this to be unnecessary in our experiments for good mouse performance.

Jointly training on both human and mouse data constrains the model slightly more than is ideal. We found that training several epochs on only one genome or the other after the full joint procedure improved validation and test set accuracy.

### GTEx SLDP

We predicted the effect of a genetic variant on various annotations by computing a forward pass through the convolutional network using the reference and alternative alleles, subtracting their difference, and summing across the sequence to obtain a single signed score for each annotation. We averaged scores computed using the forward and reverse complement sequence and small sequence shifts to the left and right. We computed scores for all 1000 Genomes SNPs, which we provide for download from [available upon publication].

Signed linkage disequilibrium profile (SLDP) regression is a technique for measuring the statistical concordance between a signed variant annotation *v* and a genome-wide association study’s marginal correlations between variants and a phenotype 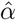 (8). The functional correlation between *v* and the true variant effects on the phenotype describes how relevant the annotation is for the phenotype’s heritability. Our model produces these signed variant annotations, and SLDP offers a validated approach to assessing their relevance to human phenotypes. Briefly, the method estimates this functional correlation using a generalized least-squares regression, accounting for the population LD structure. SLDP performs a statistical test for significance by randomly flipping the the signs of entries in *v* in large consecutive blocks to obtain a null distribution. We follow previous work in conditioning on background annotations describing minor allele frequency and binary variables for variant overlap with coding sequence (and 500 bp extension), 5’ UTR (and 500 bp extension), 3’ UTR (and 500 bp extension), and introns.

We downloaded GTEx v7a summary statistics for 48 tissues (25). We summarized each SNP’s effect on all cis-genes using the following transformation suggested for SLDP analysis

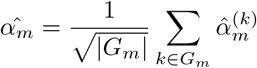

where *G*_*m*_ is the set of all genes for which a cis-eQTL test was performed for variant *m* and 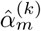 is the marginal correlation of SNP *m* and gene *k* expression (8). We passed 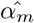 to SLDP for analysis of variant predictions.

To assess the orthogonal value of prediction scores derived from mouse datasets relative to those from human, we added the human dataset predictions to the background annotation set. Conditioning on thousands of annotations was computationally intractable. Instead, we included 64 principal components of the human variant scores matrix, which explained *>* 99% of the variance in all cases studied. To assess statistical significance, we performed the Benjamini-Hochberg procedure to correct for multiple hypotheses within each GTEx tissue.

#### Simons Simplex Collection

We downloaded 255,106 de novo variants derived from whole-genome sequencing of 1902 quartet families with an autistic child from the Simons Simplex Collection from the supplement of (29). We filtered these variants for SNPs and computed predictions as described above.

## Acknowledgments

Marta Mele-Messeguer for conversations to conceive the research direction. Jacob Kimmel, Leland Taylor, Geoffrey Fudenberg, Vikram Agarwal, and Han Yuan for valuable manuscript feedback.

## Supplemental Information

**Supplementary Figure .1:**
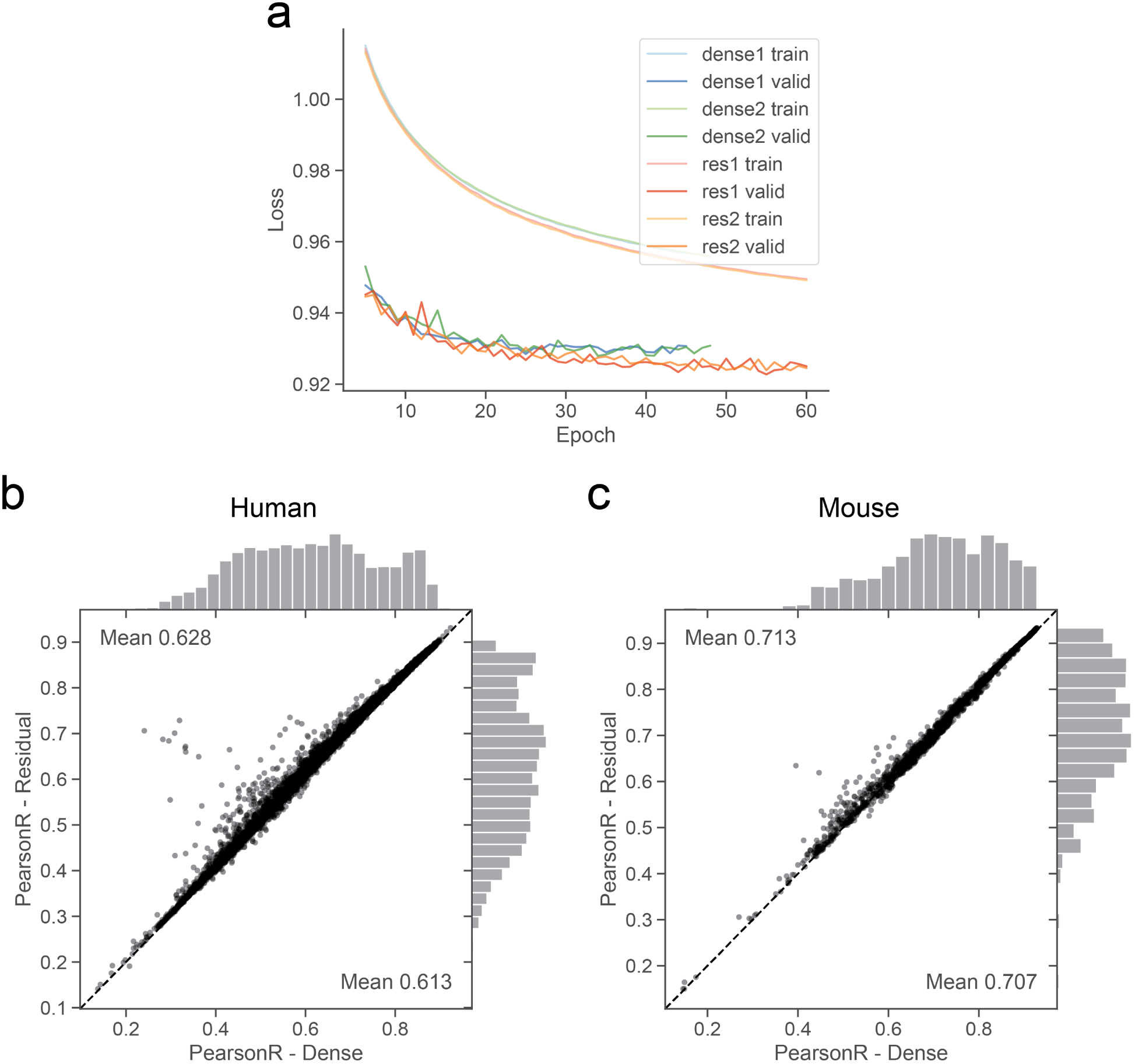
Dilated residual layers improve accuracy. We trained two separate models with approximately matched parameter totals on both human and mouse data jointly. The two models differ in how their dilated convolution layers are connected. In the first model *Dense*, which achieved the previous best for these data in (5), each layer takes all previous dilated layers as input, as opposed to taking only the preceding layer. In the second model *Residual*, introduced here, each layer takes only the previous layer as input, transforms it, and adds the new representation into the input before passing on. For each model, we computed the Pearson correlation of test set predictions and observed experimental data for thousands of datasets from various experiment types. (a) Training and validation loss curves for two replicates of the two architectures with random initializations and shuffled training examples. The top curves represent the training set and the bottom curves represent the validation set. Validation losses are less than training losses due only to stochasticity in the sequence splitting procedure. (b) Human and (c) mouse test set Pearson correlation for the best *Dense* versus *Residual* model.

**Supplementary Figure .2:**
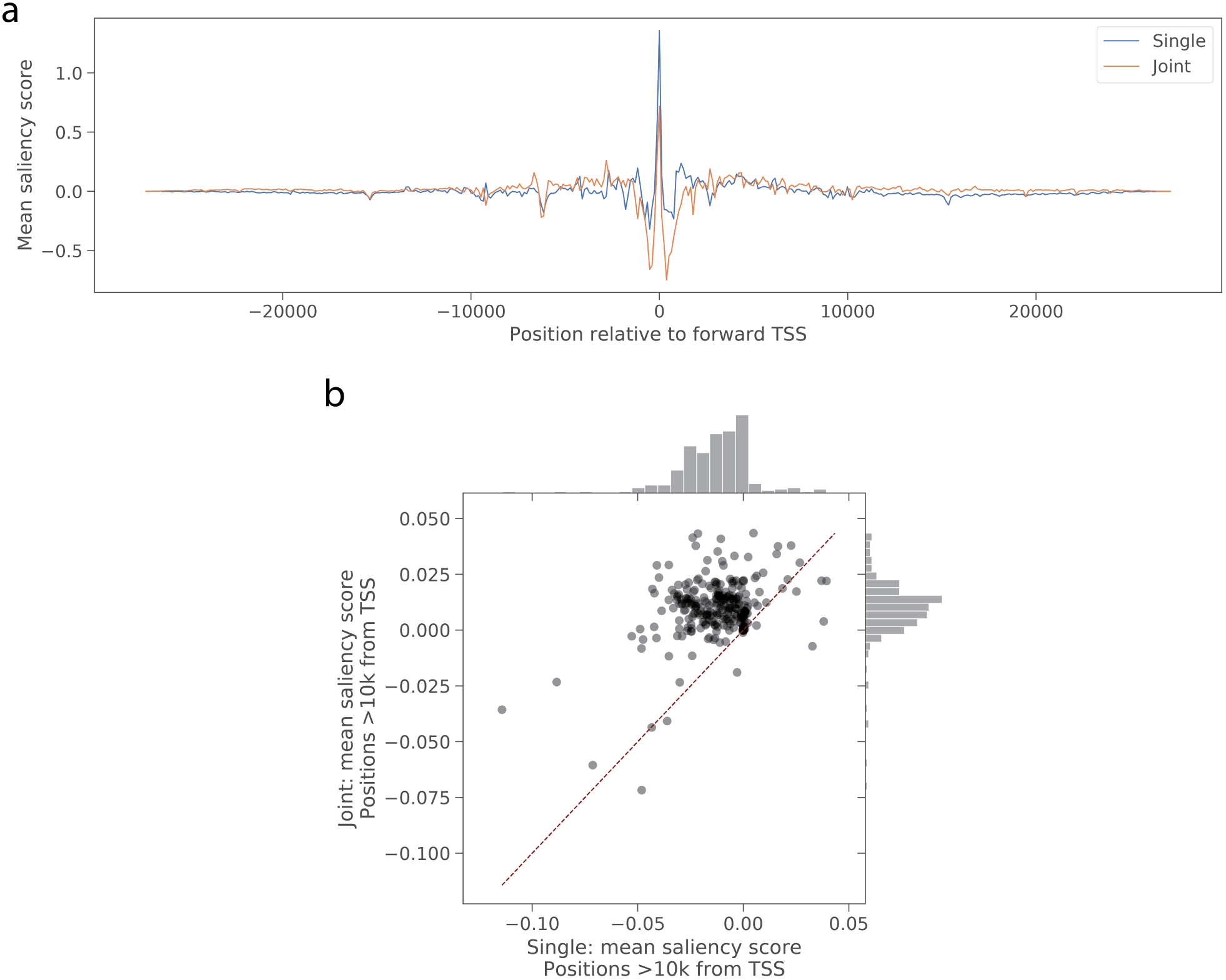
Multi-genome model assigns greater saliency scores to distal TSS regions. For 3,523 gene transcription start sites (TSS) that were not included in the training set, we computed saliency maps for the surrounding region using the model trained jointly on human and mouse (joint) and the model trained on human alone (single). The saliency map scores annotate 128-bp segments with a function of the model predictions’ gradient with respect to that segment’s vector representation after the convolutional layers and before the dilated convolutions share information across wider distances (5). Peaks in this saliency score detect distal regulatory elements, and its sign indicates enhancing (+) versus repressing () influence. (a) For each 128-bp segment, we computed the mean score across genes for liver CAGE. Patterns were consistent across CAGE datasets. (b) For segments greater than 10 kb from the TSS, the mean multi-genome model scores are greater than their single genome counterparts. This suggests that distal enhancer elements are more effectively used to predict gene expression.

**Supplementary Figure .3:**
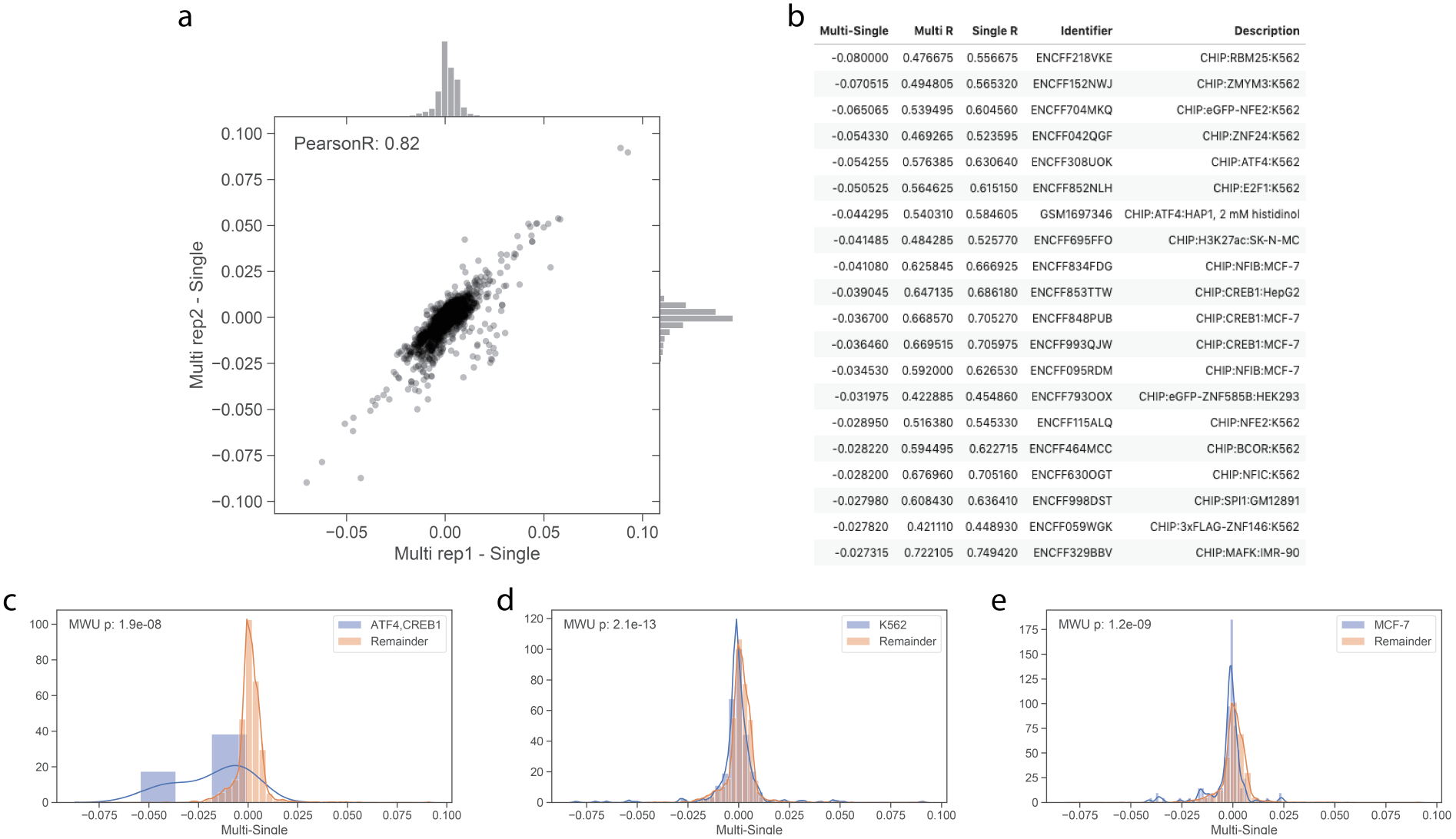
Multi-genome training harms generalization accuracy for some human ChIP-seq datasets. To further explore human ChIP-seq datasets that performed worse during multi-genome training relative to single genome training, we trained two independent replicates for both training modes. For the single genome training, we took the average of the two replicates. (a) For the multiple genome training, we plot the ChIP-seq test set PearsonR for replicate 1 minus the single genome PearsonR versus that for replicate 2. Dataset accuracy was consistent across replicates, and some ChIP-seq datasets consistently achieved lower accuracy after multi-genome training. (b) The table displays the 20 datasets with the largest decrease in test set accuracy after multi-genome training. Datasets describing (a) ATF4/CREB binding (known co-factors), (b) K562 cells, and (c) MCF-7 cells performed significantly worse according to Mann-Whitney U comparisons of the sets of 16, 476, and 129 datasets respectively. Histograms consider the average of test set PearsonR for multi-genome training minus the average for single genome training.

**Supplementary Figure .4:**
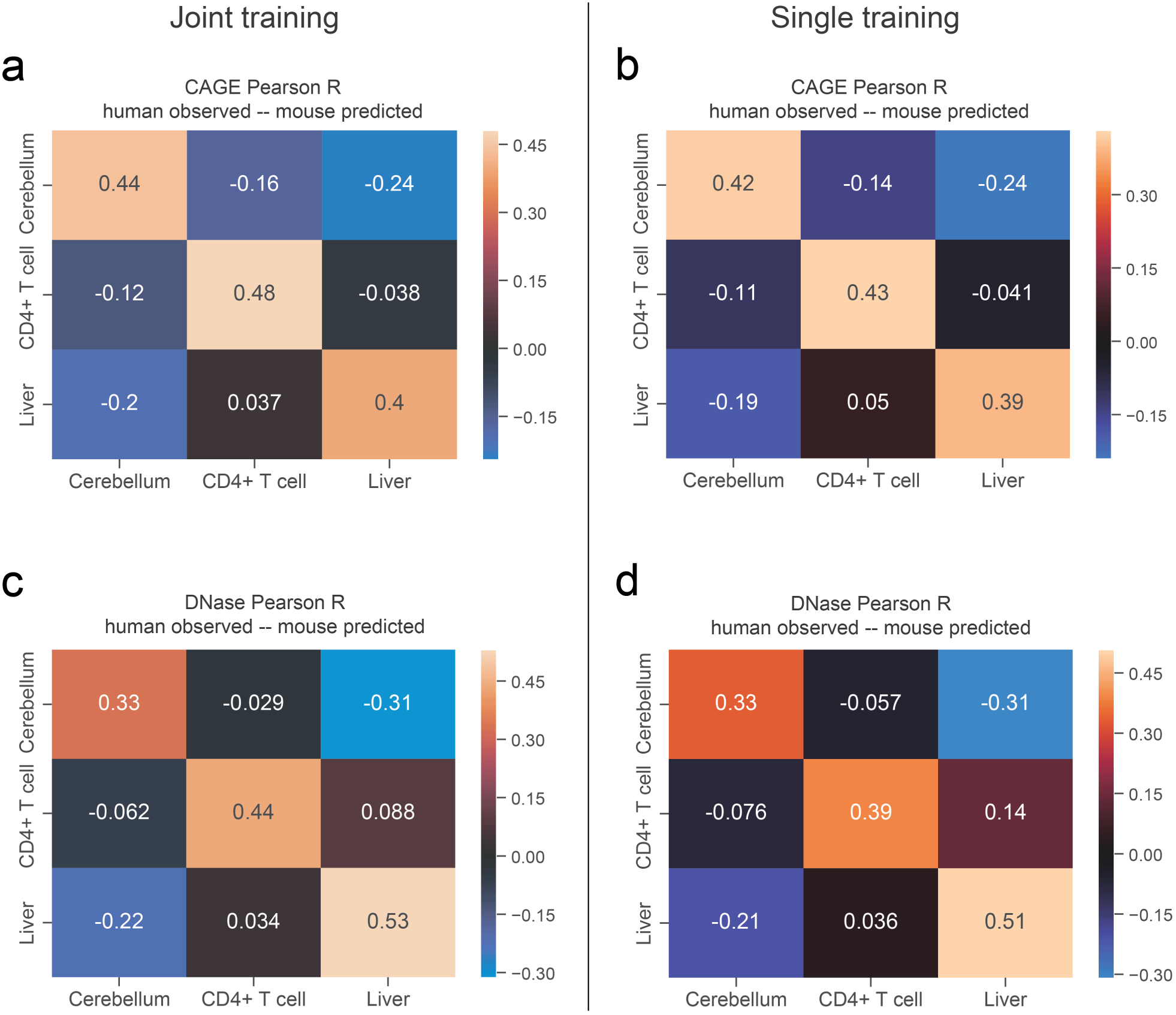
Cross-species prediction results are consistent across models. Tissue-specific regulatory programs can be learned and transferred across species, exemplified here by mouse predictions for CAGE (top row) and DNase (bottom row) for cerebellum, liver, and CD4+ T cells. Mouse predictions correspond to mouse datasets matched and compared to human datasets For CAGE, we considered the top 50% most variable TSSs, where data or predictions were quantile normalized to align sample distributions, log transformed, and mean-normalized across samples. For DNase, we considered the top 10% most variable genomic sites (less than CAGE because we consider the whole genome rather than TSSs), where data or predictions were similarly were quantile normalized to align sample distributions and mean-normalized across samples. The statistical trends were robust to most variable threshold choice. Tissue-specific cross-species accuracy depends only slightly whether the mouse model was trained jointly with human data (left column) or alone (right column). This is expected, given that the multi-genome model is more accurate on held out sequences (Figure 2).

**Supplementary Figure .5:**
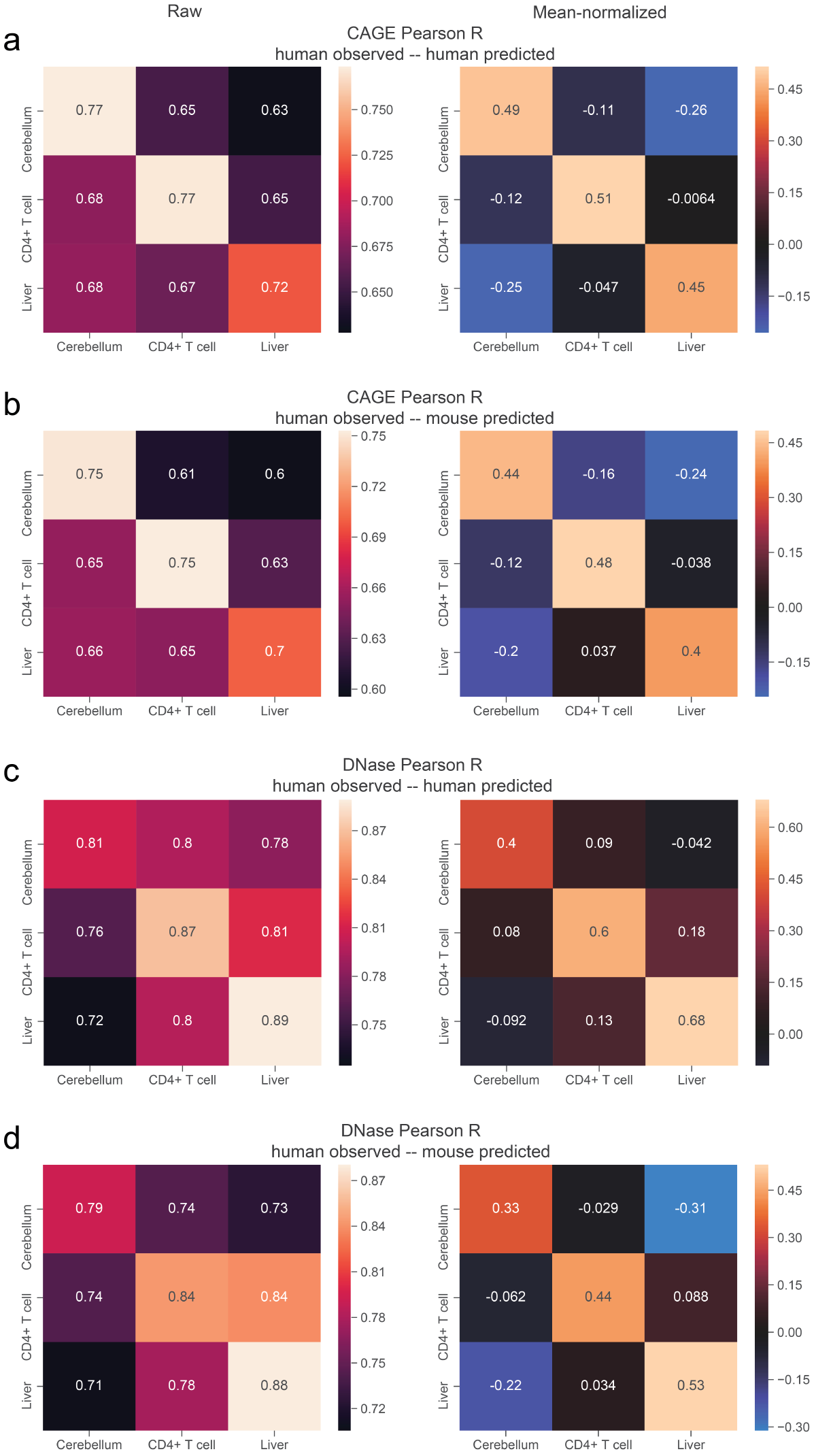
Cross-species prediction accuracy approaches that of human. Tissue-specific regulatory programs can be learned and transferred across species, exemplified here by mouse predictions for CAGE (a,b) and DNase (c,d) for cerebellum, liver, and CD4+ T cells. “Human predicted” corresponds to predictions for the human datasets, referred to as “human observed”; “mouse predicted” corresponds to predictions for the matched mouse dataset. For CAGE, we considered the top 50% most variable TSSs, where data or predictions were quantile normalized to align sample distributions, and log transformed. In the right column, we mean-normalized across samples; in the left, we did not. For DNase, we considered the top 10% most variable genomic sites (less than CAGE because we consider the whole genome rather than TSSs), where data or predictions were similarly quantile normalized to align sample distributions and mean-normalized across samples in the right column only. The statistical trends were robust to most variable threshold choice. (a,c) Human prediction accuracies exceed (b,d) mouse prediction accuracies for both CAGE and DNase.

**Supplementary Figure .6:**
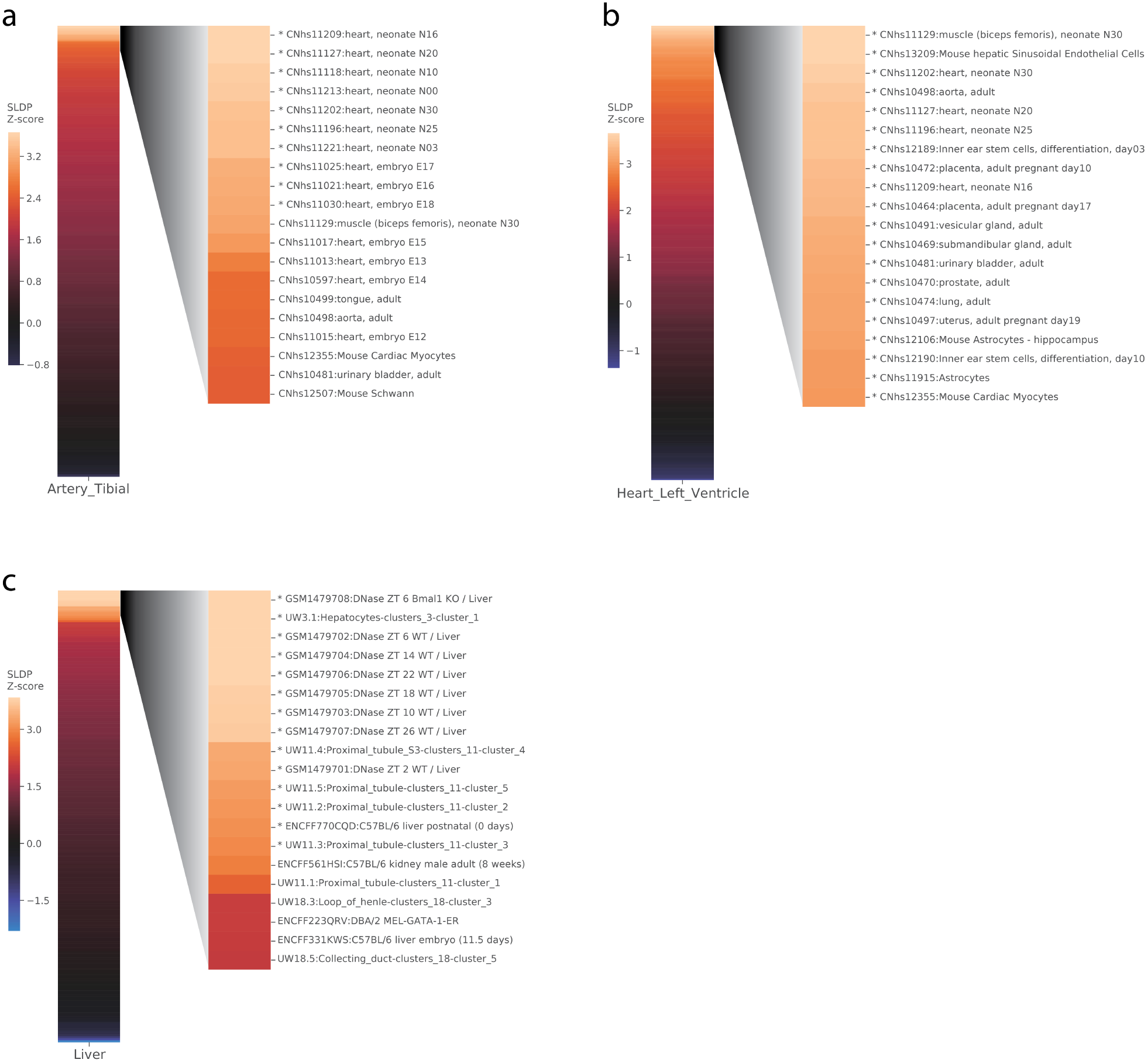
Variant effect predictions for mouse datasets significantly correlate with GTEx via SLDP, even conditional on human dataset predictions. We computed variant effect predictions for all 1000 Genomes variants with respect to human and mouse datasets. We added the first 64 principal components (PCs) of the variants by human predictions matrix, which explained 99.9% of the variance for CAGE datasets and 99.3% for DNase/ATAC. We then computed the statistical correlation between mouse predictions and GTEx summary statistics across 48 tissues using SLDP conditioned on the 64 human PCs (Methods). (a) For the tibial artery and (c) left ventricle GTEx summary statistics, mouse CAGE datasets describing the developing heart in neonate and embryo stages emerged as significant after Benjamini–Hochberg correction for multiple hypotheses. Prefix asteriks indicate FDR *q <* 0.05. Additional datasets describing adult heart components and muscle also reach significance. (b) For liver GTEx, mouse single cell hepatocyte and DNase datasets describing a 24 hour time course to profile circadian rhythms of genome accessibility reach significance.

**Supplementary Figure .7:**
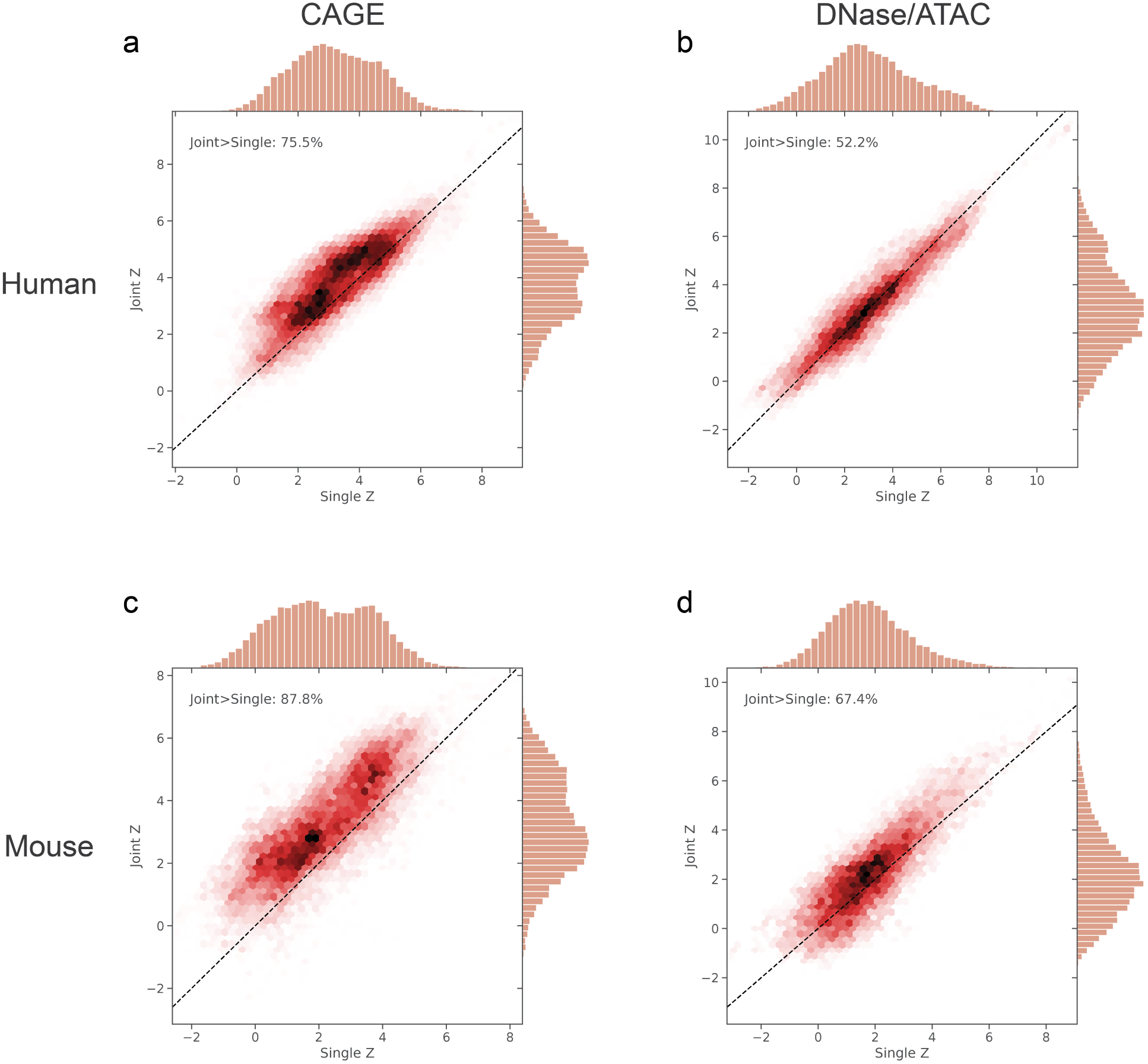
Variant effect predictions from jointly trained models correlate better with GTEx via SLDP. We computed variant effect predictions for all 1000 Genomes variants with respect to human and mouse datasets using models trained jointly on both human and mouse or trained alone on a single genome. We then computed the statistical correlation between these predictions and GTEx summary statistics across 48 tissues using SLDP (Methods). The points underlying the density maps represent every pair of model prediction dataset and GTEx tissue. SLDP signed Z-scores indicate the expected positive statistical relationship between predictions for CAGE, DNase, and ATAC-seq and gene expression. These Z-scores are clearly greater for predictions from jointly trained models for (a) human CAGE, (c) mouse CAGE, and (d) mouse DNAase/ATAC. (b) Human DNase/ATAC Z-scores are more similar between the joint and single trained models, in line with their comparable accuracy on held out sequences (Figure 2).

**Supplementary Figure .8:**
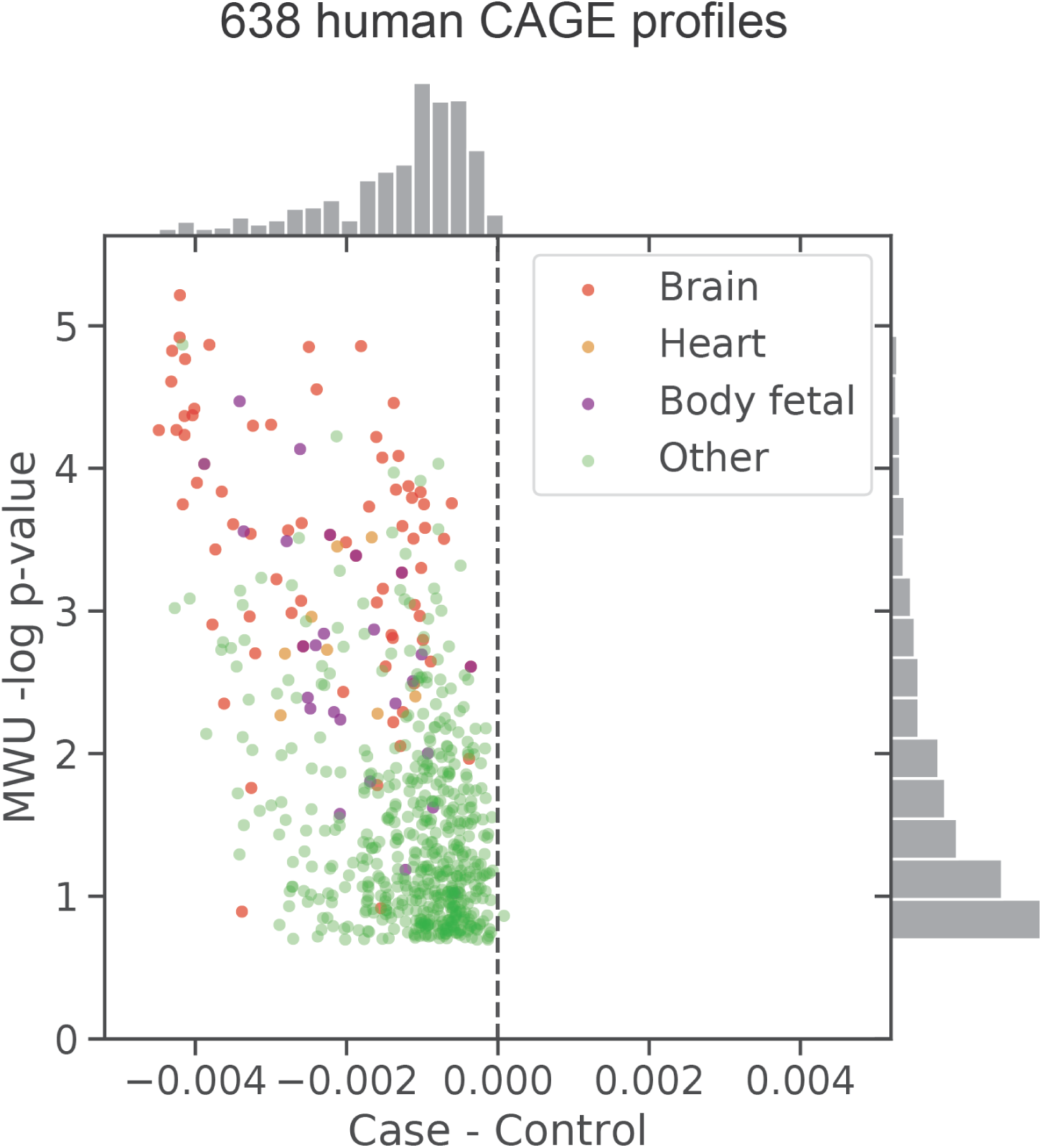
Human de novo variant predictions are more negative in the autism case versus control variants. (a) We predicted the influence of 234k de novo variants split between cases and controls on 638 CAGE datasets in human. For each dataset, we computed a Mann-Whitney U (MWU) test between case and control sets. Similarly to mouse, predictions for many datasets were enriched for more negative values in the cases, driven largely by brain, heart, and fetal tissue profiles. Correcting for multiple hypotheses using the Benjamini-Hochberg procedure among all CAGE data yielded no FDR q-values *<* 0.2. However, correcting among brain-specific CAGE data yielded 63 datasets with FDR q-value *<* 0.1. Each dataset’s x-axis position is the mean inverse hyperbolic sine over case variants minus the equivalent over control variants. The mean inverse hyperbolic sine transformation is similar to logarithm, but gracefully performs the symmetric transformation for negative values.

